# Photo-thermal Bidirectional Coupling Model for Transcranial Photobiomodulation

**DOI:** 10.64898/2026.07.11.737905

**Authors:** Ting Zhang, Yutao Chen, Xin Zeng, Ge Zhang, Feiyan Wang, Daqing Guo, Dezhong Yao

## Abstract

**Background:** Accurate optical-field simulation is critical for precise dosage delivery in transcranial photobiomodulation (tPBM). Current simulations neglect photon absorption-induced tissue heating which leads to temperature-dependent alterations of the optical field, thus fails to account for the bidirectional photo-thermal coupling effect.

**Objective:** This paper aims to establish a dynamic photo-thermal couple model that rigorously quantifies the bidirectional interaction between tissue heating and light propagation, and improves the optical dose prediction.

**Model:** We propose a Photo-Thermal bidirectional coupling Model (PTM). First, the Pennes Bioheat Equation (PBE) is employed to model the thermal response induced by photon absorption. Second, a Real-time temperature-dependent Absorption coefficient Model (RAM) is newly developed to quantify the thermal effect. Third, the PBE and RAM are integrated into the photon diffusion equation, forming the PTM. Finally, this coupled framework is solved temporally by an unconditionally stable Crank-Nicolson scheme.

**Simulations:** Benchmarking against an analytical solution (two-layer cylindrical domain) demonstrates that PTM reduces the temperature prediction error by over 2.35% compared to the uncoupled baseline. Simulations using a realistic head model reveal that, compared to PTM, uncoupled modeling underestimates energy deposition by 150 J/m^3^ and overestimates photon fluence by 3 J/m^2^ within just one minute, and such discrepancies will be amplified with increasing exposure duration and power. Furthermore, the PTM identifies a 1.43-mm advantage in penetration depth for pulsed-wave over continuous-wave modality under iso-energy conditions, a key insight enabled by the coupled modeling approach.

**Conclusion:** The PTM provides a high-fidelity simulation framework that captures the dynamic, bidirectional photo-thermal coupling in tPBM, explicitly quantifying the thermal feedback ignored by current models and thereby enabling more reliable treatment optimization and safer clinical translation.

## 1. Introduction

Transcranial photobiomodulation (tPBM) has emerged as a promising non-invasive neuromodulation modality, demonstrating potential for treating neurological disorders and enhancing cognitive functions [1, 2, 3]. Its primary therapeutic mechanism involves the resonant activation of cytochrome C oxidase (CCO) within the mitochondrial respiratory chain, subsequently up-regulating adenosine triphosphate (ATP) production and cerebral energy metabolism [4, 5, 6]. However, the clinical translation of tPBM is significantly hampered by high individual variability in treatment outcomes [7, 8, 9]. This variability stems primarily from two factors: (1) subject-specific anatomical heterogeneity, which deflects light propagation [10]; and (2) the well-documented biphasic dose-response nature of tPBM, where excessive energy yields inhibitory rather than beneficial effects [11, 12]. Consequently, developing high-fidelity, personalized computational optical field model to accurately predict the treatment dosage is imperative for precision neuromodulation.

Existing modeling approaches for tPBM light propagation generally follow two directions. The first prioritizes computational efficiency, utilizing analytical solutions for simplified geometries, such as cylinders and spheres [13]; however, these methods struggle to capture light migration in the complex head geometry. The second direction emphasizes prediction accuracy through sophisticated numerical methods, including Monte Carlo simulations (e.g., MMC [14], MCVM [15], MCML [16]) and finite element analysis (e.g., NIRFAST [17]). These advanced approaches are highly adept at accommodating the anatomically realistic, heterogeneous structures of the human head, as well as modeling complex wide-field illumination schemes. Furthermore, the advent of GPU-accelerated and mesh-based Monte Carlo (MMC) algorithms has profoundly elevated both the computational efficiency and spatial precision of these simulations [18, 19, 20]. Despite these substantial methodological improvements, a fundamental limitation persists: these models universally operate under the assumption that absorbed photon energy is either consumed photochemically or permanently lost from the head system, completely neglecting its conversion to heat and the consequential thermal feedback on light distribution.

While tPBM is primarily driven by photochemical mechanisms, a non-negligible fraction of energy is dissipated as heat via non-radiative relaxation, resulting in localized temperature elevations [21]. Experimental measurements corroborate this phenomenon, with skin absolute temperatures reaching up to 40 °C under typical tPBM parameters [22]. Importantly, beyond primary photochemical pathways, emerging evidence suggests that such localized heating may play a secondary role in neuromodulation through the potential activation of heat-sensitive targets or the enhancement of local microcirculation. Consequently, researchers have begun to characterize these thermal dynamics using the Pennes Bioheat Equation (PBE) [23, 24], a framework originally established to assess electromagnetic-induced heating and optimize photothermal tumor ablation [25]. However, by treating heat generation merely as a static, terminal output, these established models completely ignore the reciprocal feedback of temperature-dependent physicochemical changes on tissue optical properties, presenting a fundamental barrier to accurate dosimetry.

From a thermodynamic perspective, the dissipated thermal energy actively modifies the field distribution by modulating the optical properties of biological tissues, principally the absorption coefficient. Experimental studies demonstrate a significant temperature dependence of the absorption coefficient in skin tissue, largely attributable to the strong temperature sensitivity of moisture content in the near-infrared spectrum [26]. For example, water absorption increases by approximately 10% as temperature rises from 20 °C to 40 °C, accompanied by a blue shift in the absorption peak (from 971 nm to 966 nm) [17]. Furthermore, numerous investigations have confirmed an approximately linear positive correlation between the absorption coefficient and temperature within physiological limits (up to 42 °C) for both human skin and brain [28, 29, 30]. In light of this, we believe that the thermal feedback to the optical field should be considered by establishing a Real-time temperature-dependent Absorption coefficient Model (RAM).

Building upon these theoretical basis and experimental facts, this paper will introduce a Photo-Thermal bidirectional coupling Model (PTM) that integrates the PBE with a temperature-dependent constitutive model for optical absorption, thereby formalizing the coupled spatiotemporal dynamics. The principal contributions of this work are threefold:

- A novel coupled computational framework (PTM) is proposed to enhance the simulation accuracy of both optical and thermal fields in tPBM.
- A thermal feedback model (RAM) is formulated and embedded into the PTM to dynamically quantify the thermal effect on light propagation.
- The proposed PTM enables spatiotemporal tracking of photo-thermal dynamics in tPBM, facilitating the investigation of diverse stimulation modalities.

The paper is organized as follows: Section 2 presents the governing equations of the PTM; Section 3 describes its numerical implementation, evaluation metrics, and the design of computational experiments; Section 4 details the experimental results; Section 5 discusses findings and future work; finally, Section 6 concludes this paper.

## 2. The photo-thermal bidirectional coupling model

This section first introduces the mathematical-physical equations that govern photon propagation and heat conduction in the head domain. Then, the theoretical foundations of bidirectional coupled modeling are established. The overview of the coupling framework is illustrated in Fig. 1.

**Fig. 1.**
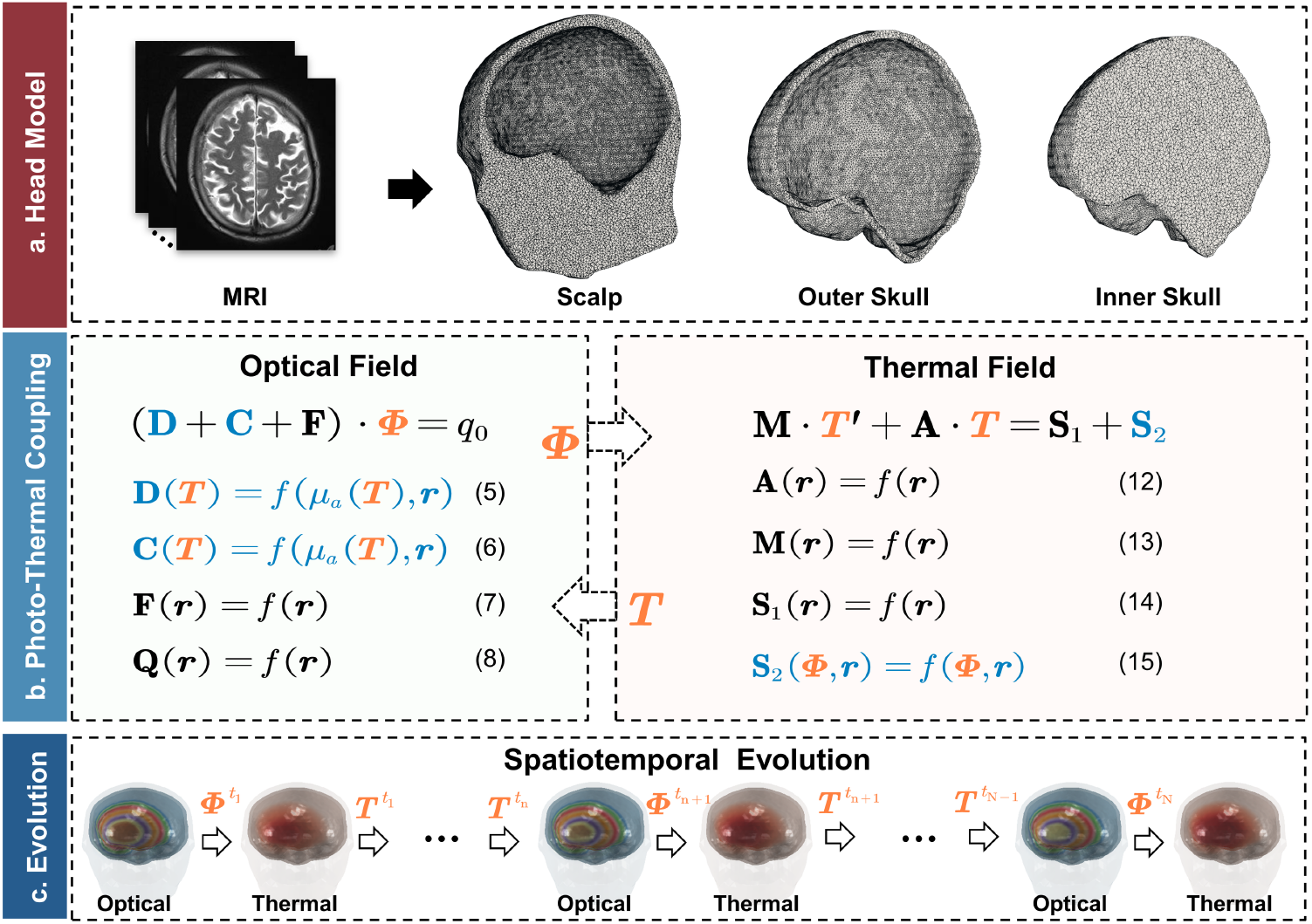
An overview of the proposed Photo-thermal bi-directional coupled modeling framework. (a) Head model construction: Based on the 6th-generation ICBM 152 (International Consortium for Brain Mapping 152) nonlinear head template, a three-layer head model (scalp, outer skull, inner skull) consisting of 515,704 nodes and 3,088,914 elements is built for photo-thermal simulation. (b) Photo-thermal bidirectional coupled model: The left and right panels depict the discretized governing equations for the optical field (Eqs. (5) – (8) in the main text) and the thermal field (Eqs. (12) – (15) in the main text), respectively. Blue-marked terms represent the temperature - dependent optical properties and the light-intensity-dependent heat sources, while orange - marked terms denote the bidirectionally coupled variables (photon flux ***Φ***, temperature ***T***). (c) Schematic diagram illustrating the spatiotemporal evolution of the optical and thermal fields.

### 2.1. Mathematic-Physical Equation

#### 2.1.1. Diffusion Approximation of tPBM

An optical field is considered “diffuse” when the isotropic fluence significantly exceeds the directional flux. This condition arises in scattering-dominated regimes, far from sources and boundaries, provided that the fluence does not vary rapidly in time (e.g., on sub-picosecond scales). This assumption enables the simplification from the general radiative transport equation, which describes anisotropic fields, to the diffusion approximation, suited for isotropic fluence [17, 31]. The diffusion approximation in the frequency domain is given by

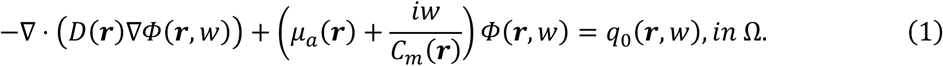

Here, Ω is the solution domain of the target light field, i.e., head tissue; *Φ*(***r***, *w*) is the photon flux at position ***r***, and modulation frequency *w*, with unit 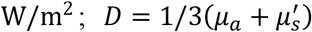 is the diffusion coefficient; *μ*_*a*_ and 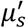 are the absorption and reduced scattering coefficients, respectively; *C*_*m*_(***r***) is the light speed in the biological tissue at position ***r***; *q*_0_(***r***, *w*) is the isolated source at position ***r***, and modulation frequency *w*.

An index-mismatched Robin boundary condition [32, 33] was applied at the external boundary. This condition prescribes that the outgoing flux is equal to the local flux multiplied by a factor that models the internal reflection, preventing any backflow of light from the air into the tissue. This relationship is defined by the following equation:

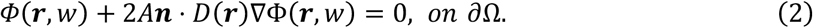

Here, ∂Ω is the external surface of Ω and ***r*** is a point on ∂Ω; ***n*** is the outward-pointing normal; *A* depends on the refractive index (RI) mismatch between scalp and air. Here, *A* can be derived from Fresnel’s law:

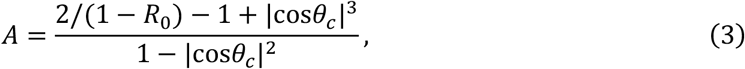

where *θ*_*c*_ = arcsin(*n*_*air*_/*n*_1_) is the critical angle for total internal reflection of photons transitioning from region Ω with RI *n*_1_ to the external air with RI *n*_*air*_, and *R*_0_ = (*n*_1_/*n*_*air*_ − 1)^2^/(*n*_1_/*n*_*air*_ + 1)^2^ is the normal incidence reflectivity at the boundary. Conventionally, the external boundary RI is assumed equal to that of free space, i.e., *n*_*air*_ = 1.

To solve equation (1) numerically, the weak form and a discretization scheme are applied. Readers are referred to the Supplementary Material for the detailed derivations. This process generates an *N* × *N* linear system that is used to compute the element stiffness matrices and assemble the global stiffness matrix, as shown below:

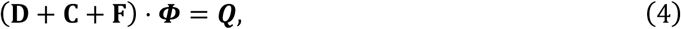

where **D, C, F** ∈ ℝ^*N*×*N*^ are the terms related to the geometry and the physical properties of tissues; **Q** ∈ ℝ^*N*×1^ is a vector related to light sources; ***Φ*** = (*Φ, Φ*_2_, …, *Φ*_*N*_) ∈ ℝ^*N*×1^ is the numerical solution obtained from this system, which serves as the desired approximation to the photon flux; and Ω_*h*_ represent the discrete domain and *N* the number of mesh nodes. **D, C, F** and **Q** are given by:

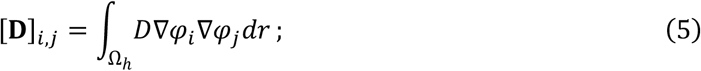

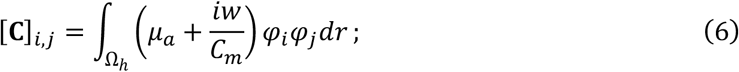

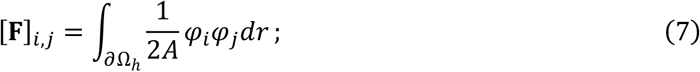

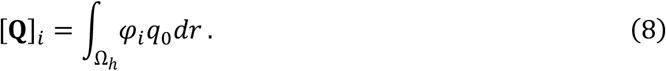

Here, *φ*_*i*_ and *φ*_*j*_ are piecewise linear global shape functions associated with nodes *i* and *j* on the discrete three-dimensional domain Ω_*h*_.

#### 2.1.2. Pennes Bioheat Equation

The energy conservation for biological heat transfer is governed by the PBE, derived from Fourier’s law of heat conduction. Thus, the PBE enables the direct calculation of temperature changes resulting from light absorption across tissue layers.

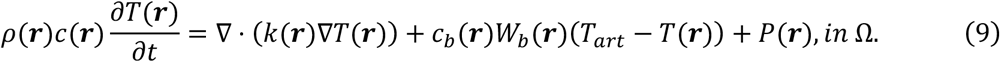

Here, Ω is the solution domain of the target thermal field, i.e., head tissue; *T*(***r***) is the temperature at position ***r***; *ρ*(***r***), *c*(***r***) and *k*(***r***) represent the tissue density, specific heat, and thermal conductivity at position ***r*** respectively; *T*_*art*_ signifies constant arterial temperature (set at 37 °C in this paper); *W*_*b*_(***r***) and *c*_*b*_(***r***) are the perfusion rate and specific heat of blood at position ***r***; *c*_*b*_(***r***)*W*_*b*_(***r***)(*T*_*art*_ − *T*(***r***)) represents the thermal energy contribution from the blood perfusion; *P*(***r***) describes the energy received from external sources at position ***r***, unit: W/m^3^.

Given the heat transfer between the scalp and the external environment, a natural convection (Robin) boundary condition is applied.

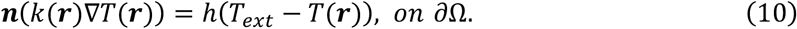

Here, ∂Ω is the external surface of Ω and ***r*** is a point on ∂Ω; ***n*** is the outward-pointing normal; *h* is the heat transfer coefficient, assigned a value of 6.2 W*m*^−2^K^−1^ [34], and *T*_*ext*_ is the ambient temperature, set to 25 °C.

The finite element approximation to the above PBE leads to a system of ordinary differential equations (ODEs). The complete derivation refers to the Supplementary Material.

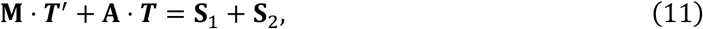

where **M, A** ∈ ℝ^*N*×*N*^ denote the mass and stiffness matrices respectively; the vectors **S**_1_, **S**_2_ ∈ ℝ^*N*×1^ represent source terms, with **S**_1_ accounting for the heat exchange with blood and the environment, and **S**_2_ characterizing the light absorption; and ***T*** = (*T*,, *T*_2_, …, *T*_*N*_) ∈ ℝ^*N*×1^ is a temperature vector of the discrete solution (Ω_*h*_). The explicit expression for **M, A, S**_**1**_ and **S**_**2**_ are defined as follows:

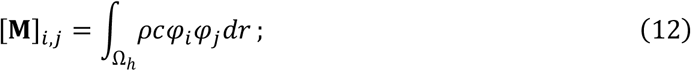

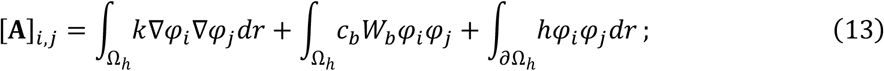

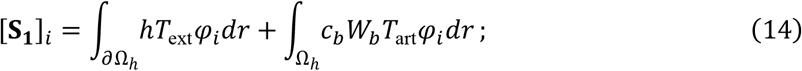

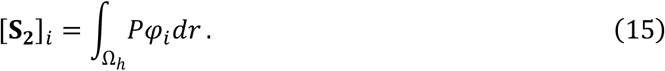

Here, *φ*_*i*_ and *φ*_*j*_ are piecewise linear global shape functions associated with nodes *i* and *j* on the discrete three-dimensional domain Ω_*h*_. Furthermore, this ODEs system is discretized in time using the Crank-Nicolson method (with *θ* = 0.5). This method is chosen for its unconditional stability, which mitigates the impact of mesh size on the solution.

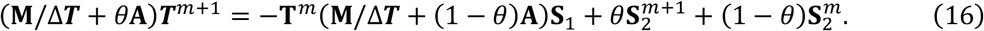

### 2.2. Bidirectional Coupled Modeling

The bidirectional coupled modeling comprises two components: (1) the light energy that penetrates biological tissue leads to changes in temperature; (2) these temperature variations modify the physical characteristics of the optical field by adjusting optical properties.

#### 2.2.1. Forward Coupling

The objective of forward coupling is to model the impact of light transport on bioheat transfer. This is achieved by incorporating the absorbed energy density obtained from equation (1), as a heat source, into the PBE, which is a well-established approach in laser thermal therapy studies [35, 36, 37]. Here, the absorbed power density is defined as *P* = *μ*_*a*_*Φ*. It is crucial to note that the light source in our model is distributed across the entire computational domain (i.e., the whole head domain), rather than being confined solely as a boundary condition for illumination.

#### 2.2.2. Back Coupling

The primary objective of the thermal feedback coupling is to establish the Real-time temperature-dependent absorption coefficient Model (RAM). The RAM is designed to characterize the temperature (*T*) dependence of the absorption coefficient (*μ*_*a*_) and mathematically formulate their quantitative relationship. Experimental studies indicate a significant positive linear correlation between *μ*_*a*_ and *T* within the relevant therapeutic range (where *T* is maintained below the ∼42°C threshold for cellular hyperthermia damage). Consequently, we suppose the relationship following the formula (17):

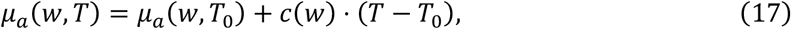

where *w* is the wave length; *T*_0_ and *T* are the reference and current temperatures, respectively; *c*(*w*) = ∂*μ*_*a*_(*w, T*)/ ∂*T* is the temperature sensitivity coefficient of the absorption coefficient *μ*_*a*_.

## 3. Simulation Method

### 3.1. MRI Data and Head Model

#### 3.1.1. MRI Data

The anatomical basis for the simulations was established using the 6th-generation “ICBM 152” nonlinear template. Sourced from the Brainstorm software distribution, this T1-weighted MRI template represents a population average of 152 healthy human brain scans from the Montreal Neurological Institute (MNI) database [38, 39, 40]. Based on this template, a three-dimensional head model was constructed to serve as the unified computational domain for both the optical and thermal simulations.

#### 3.1.2. Head Model

The construction of the computational head model involved generating a labeled volumetric mesh and defining the optical incident site. First, using the Iso2mesh toolkit [41], we generated a three-dimensional tetrahedral mesh comprising 515,704 nodes and 3,088,914 elements. This anatomical model was segmented into three distinct tissue compartments: scalp, skull, and brain. The wavelength-specific optical properties (at 810 nm) and thermal characteristics assigned to each layer are summarized in Table 1. Subsequently, to target the right prefrontal cortex, the simulated optical source was spatially co-registered to the “Fp2” position of the international 10-10 EEG system, using the interactive alignment module within the FieldTrip toolbox [42].

**Table 1.** Tissue types and optical and thermal properties [43, 44, 45].

| Parameter | Scalp | Skull | Brain |
| --- | --- | --- | --- |
| $\mu_a$ (1/mm) | 0.0450 | 0.0110 | 0.0600 |
| $\mu'_s$ (1/mm) | 2.1800 | 1.9200 | 2.8715 |
| $\partial\mu_a/\partial T$ (mm<sup>-1</sup> · K<sup>-1</sup>) | $1.0922 \times 10^{-5}$ | 0 | $1.865 \times 10^{-6}$ |
| $\rho$ (kgm<sup>-3</sup>) | 1070 | 1520 | 1030 |
| $c$ (J · K<sup>-1</sup>kg<sup>-1</sup>) | 3662 | 1590 | 3640 |
| $k$ (W · K<sup>-1</sup>m<sup>-1</sup>) | 0.3420 | 0.6500 | 0.5340 |
| $c_b$ (J · K<sup>-1</sup>kg<sup>-1</sup>) | 3900 | 3900 | 3900 |
| $W_b$ (kg · m<sup>-3</sup>s<sup>-1</sup>) | 1.8000 | 0.0600 | 6.9500 |

#### 3.1.3. Thermal Sensitivity Coefficient

To formalize the thermal feedback within the RAM, the temperature sensitivity coefficients of the absorption properties (*c*(*w*) = ∂*μ*_*a*_(*w, T*)/ ∂*T*) at 810 nm were determined for the scalp, skull, and brain based on empirical data. These layer-specific coefficients were parameterized as follows: (1) for the scalp, the coefficient was analytically derived from formula (18) [46]; (2) for the skull, the coefficient was assumed to be zero, justified by its remarkably low water content and negligible photo-thermal response [47, 48]; and (3) for the brain tissue, a validated constant of 1.865 × 10^−1^ *m*^−1^K^−1^was adopted from established literature [29].

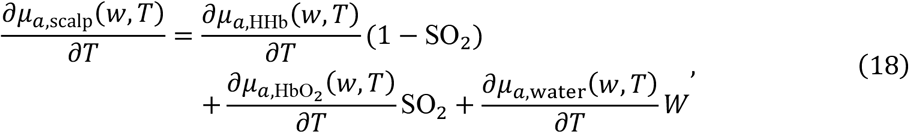

where 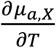 for *X* ∈ {HHb, HbO_2_, water, *scalp*} represent the thermal sensitivity coefficients of the respective chromophores or tissue. Their values are assigned as −9.6774 × 10^−6^ *mm*^−1^s^−1^ for HHb, 2.2581 × 10^−5^ *mm*^−1^s^−1^ for H*b*O_2_, and 3.8400 × 10^−6^ *mm*^−1^s^−1^ for water, based on [27, 49, 50]. SO_2_ and $*W*$ represent the oxygen saturation and water content, where the values for skin are specified as SO_2_ = 0.71 and *W* = 0.6, following [46].

### 3.2. Stimulation Settings

To systematically investigate the bidirectional photo-thermal dynamics and validate the proposed PTM, distinct optical stimulation paradigms were configured for two different computational domains, i.e., a realistic head domain and a two-layer cylindrical domain. For physiological investigations utilizing the realistic head model, clinically representative continuous-wave (CW) and pulsed-wave (PW) stimulation parameters (detailed in Table 2) were applied. Furthermore, we comprehensively modulated the irradiance and duty cycle to quantify the specific impact of these dosimetric variables on the photo-thermal coupling process. Conversely, to rigorously benchmark the numerical accuracy of the PTM, a standardized high-irradiance CW stimulation condition (1 W/*cm*^2^, 600 *s*) was configured for the two-layer cylindrical phantom. This specific parameter, though exceeding typical clinical thresholds, was intentionally selected to induce a pronounced thermal gradient, thereby facilitating a direct and robust comparison between the numerical predictions of the PTM and the analytically derived ground truth.

**Table 2.** Laser parameters used in simulation for the realistic head model.

| Laser parameters | Values |
| --- | --- |
| Wavelength (nm) | 810 |
| Laser mode | CW, PW |
| CW power density ($\text{W/cm}^2$) | 0.5, 1 |
| PW peak power density ($\text{W/cm}^2$) | 0.25, 0.5, 0.75, 1 |
| PW average power density ($\text{W/cm}^2$) | 0.1, 0.3, 0.5, 0.7, 0.9, 1 |
| Area of laser beam radiation ($\text{cm}^2$) | 12.57 |
| Pulse frequency (Hz) | 10 |
| Duty cycle | 0.1, 0.3, 0.5, 0.7, 0.9 |
| Irradiation time (s) | 60, 600 |

### 3.3. Evaluation

This paper proposes a novel PTM that dynamically incorporates temperature effects to achieve high-fidelity modeling of light propagation in the head domain during tPBM. The proposed PTM enables the simultaneous spatiotemporal computation of photon fluence distributions and thermal field evolution, thereby facilitating robust dosimetric planning and thermal safety assessments. To assess the performance of this model, we compare its numerical solutions with those of the conventional uncoupled photon model and evaluate the differences in photon and temperature distribution utilizing the relative difference measure (RDM) [51, 52].

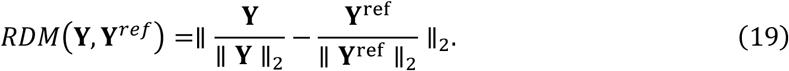

Here, ∥⋅∥_2_ denotes the Euclidean norm. **Y** is a vector of values (temperature, photon flux, energy deposition, or absorption coefficient) at the target mesh nodes for the comparison group, and **Y**^ref^ is the vector of values at the corresponding nodes for the reference group.

To systematically validate the PTM and elucidate the bidirectional photo-thermal dynamics, a progressive series of four computational experiments was designed.

#### 3.3.1. Experiment I for a two-layer cylindrical domain with analytical solution

To rigorously verify the numerical accuracy of the PTM, numerical predictions from both the proposed framework and the current uncoupled baseline are benchmarked against an exact analytical solution reported by [13, 53] for a two-layer cylindrical domain (with the explicit expression provided in Supplementary Material S3). The numerical geometry (radii: 18 and 25 mm; height: 1100 mm) and the internal source configuration (radius: 20 mm, 60° arc) are specifically tailored to satisfy the infinite-length and boundary assumptions required by the analytical method.

#### 3.3.2. Experiment II for realistic head model

This experiment quantifies the spatiotemporal thermal effects of tPBM and the resulting dynamic feedback on the temperature-modulated absorption. Systematic parametric sweeps within clinical ranges (Table 2) include: (1) six duty cycles from 10% to 90% at a constant peak power density of 1 W/*cm*^2^, (2) four pulsed-wave peak power densities from 0.25 to 1 W/*cm*^2^ at a fixed 90% duty cycle, and (3) six average power densities ranging from 0.1 to 1 W/*cm*^2^.

#### 3.3.3. Experiment III for realistic head model

To demonstrate the necessity of bidirectional coupling, the PTM is benchmarked against a current uncoupled model. We quantify discrepancies in spatiotemporal photon fluence and energy deposition by: (1) tracking temporal deviations using the parametric sets from Experiment I, and (2) mapping spatial disparities across peak irradiances from 0.25 to 1 W/*cm*^2^ at a 90% duty cycle.

#### 3.3.4. Experiment IV for realistic head model

Leveraging the dynamic spatiotemporal capability of the PTM, we conduct a comparative study between CW and PW irradiation, an analysis precluded by the static-field assumption of current models. To ensure an iso-energetic comparison, both modalities are simulated at an identical average irradiance (0.5 W/cm^2^) for 600 seconds. We systematically evaluate their resultant temperature profiles, dynamic absorption coefficients, photon fluence, energy deposition, and penetration depths to uncover how the distinct energy delivery profiles of CW and PW drive their differential therapeutic efficacies.

## 4. Simulation Results

Our simulation analysis evaluates the spatiotemporal evolution of temperature, the absorption coefficient, photon flux, photon fluence, and energy deposition, extracted from the finite element mesh. To prevent the spatial dilution of averages caused by large non-irradiated peripheral regions, data extraction is strictly confined to a cylindrical region of interest (ROI) beneath the illumination site. Set to 1.5 times the source radius and coaxially aligned with the beam, this ROI fully encapsulates the core photon and thermal diffusion regions. The subsequent results are systematically presented in three formats: (1) time series of spatial averages; (2) depth-dependent spatial profiles based on last-pulse averages (temperature and *μ*_*a*_) and the cumulative response over the entire stimulation period (photon fluence and energy deposition); and (3) spatial distribution maps of model discrepancies.

### 4.1. Comparison with Analysis Solution

As a foundational step to ensure computational reliability, we first conduct a benchmark comparison against the exact analytical solution and the numerical results from both the coupled and uncoupled models in a two-layer cylindrical domain with analytical solution. Fig. 2 presents the computational setup and results. Panel (a) details the computational domain and mesh used for finite element simulations, while panel (b) compares the temperature distributions on the inner cylinder alongside the RDMs between two numerical models and the analytical reference. Macroscopically, the color bars across all three approaches exhibit comparable dynamic ranges, indicating similar simulated thermal bounds. Quantitatively, as shown in Fig. 2b, the RDMs of both numerical models decrease exponentially with irradiation time. Compared to the uncoupled model, the coupled model maintains a persistently lower RDM, achieving a relative error reduction of 2.35% to 8.62% across the entire simulation duration. Qualitatively, the spatial distribution generated by the coupled model aligns closely with the analytical benchmark, whereas the uncoupled model exhibits more pronounced spatial discrepancies, particularly near geometric boundaries.

**Fig. 2.**
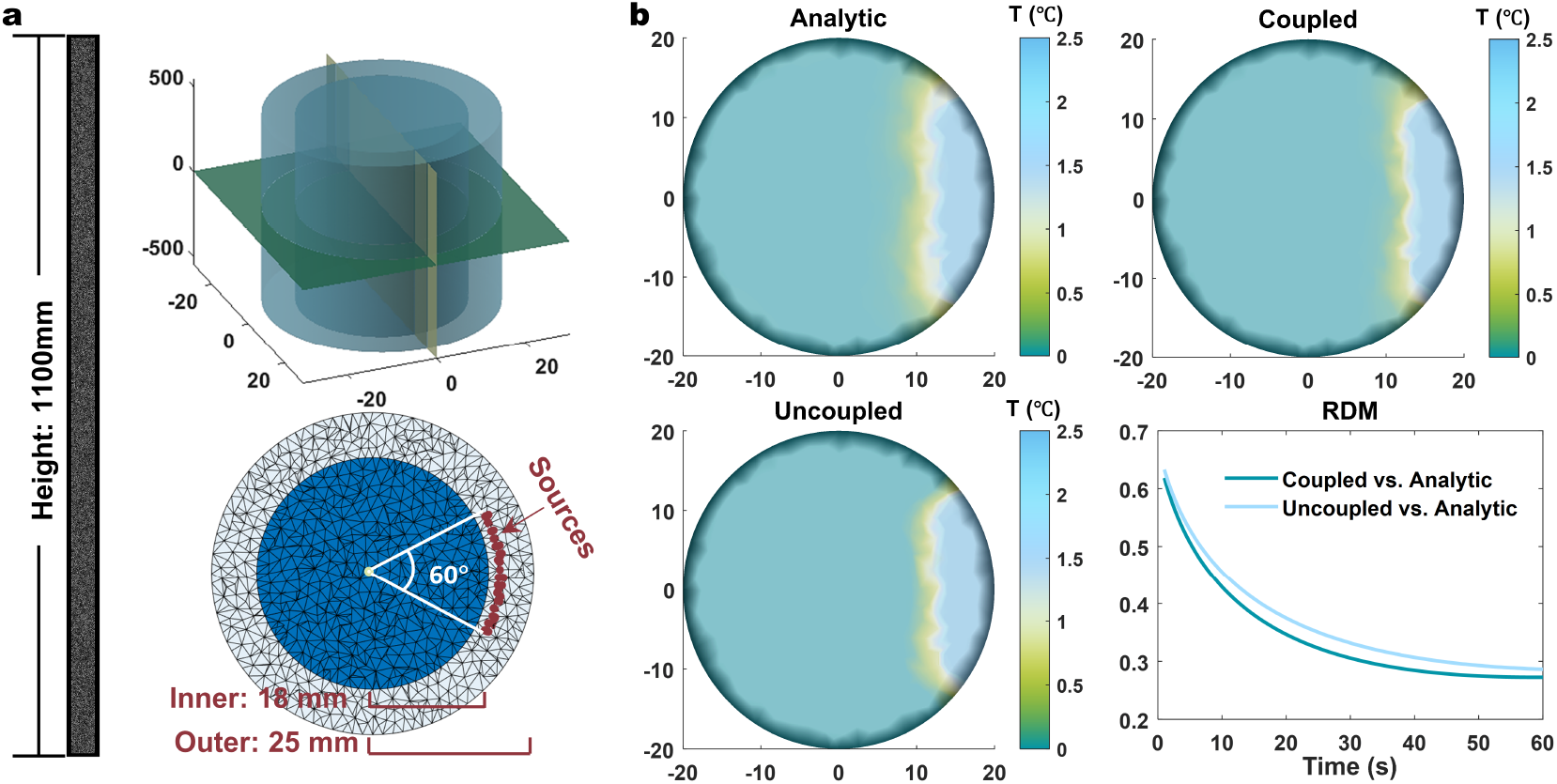
Benchmarking of numerical models (the proposed PTM and current uncoupled model) against an analytical solution for a two-layer cylindrical domain effectively infinite cylinder. (a) Computational domain and mesh. (b) Comparison of radial distributions and the relative difference measure.

### 4.2. Thermal Accumulation and Temperature-Mediated Optical Regulation

The thermal accumulation in tPBM is presented in Fig. 3 in terms of parameter sensitivity and spatiotemporal dynamics in the realistic head model. At the parameter level, it is observed from Fig. 3a illustrates a progressive temperature elevation from below 37°C to approximately 37.4°C within one minute. Specifically, increasing the duty cycle from 10% to 100% amplifies the average temperature change from less than 0.2°C to over 0.4°C (see the detailed view). Similarly, as the peak power density (P_peak_) scales from 0.25 to 1 W/cm^2^, the transient temperature rise grows proportionally (Fig. 3b). Spatially, this thermal elevation is predominantly confined to superficial layers (about 0–10 mm) and decays exponentially with depth, with a peak increase at the surface of less than 2 °C after one minute (Fig. 3c). Temporally, the active heating phases are tightly synchronized with the pulse-on periods, interspersed with complex off-time dynamics where the temperature briefly continues to rise before transitioning into a stable plateau (Figs. 3a, b). Tracking these thermal dynamics, *μ*_*a*_ exhibits commensurate temporal variations. Under elevated duty cycle or P_peak_, *μ*_*a*_ increases by approximately 6 × 10^−4^m^-1^ within one minute, with the maximum superficial variations remaining below 0.02 m^-1^ (Fig. 3e, f).

**Fig. 3.**
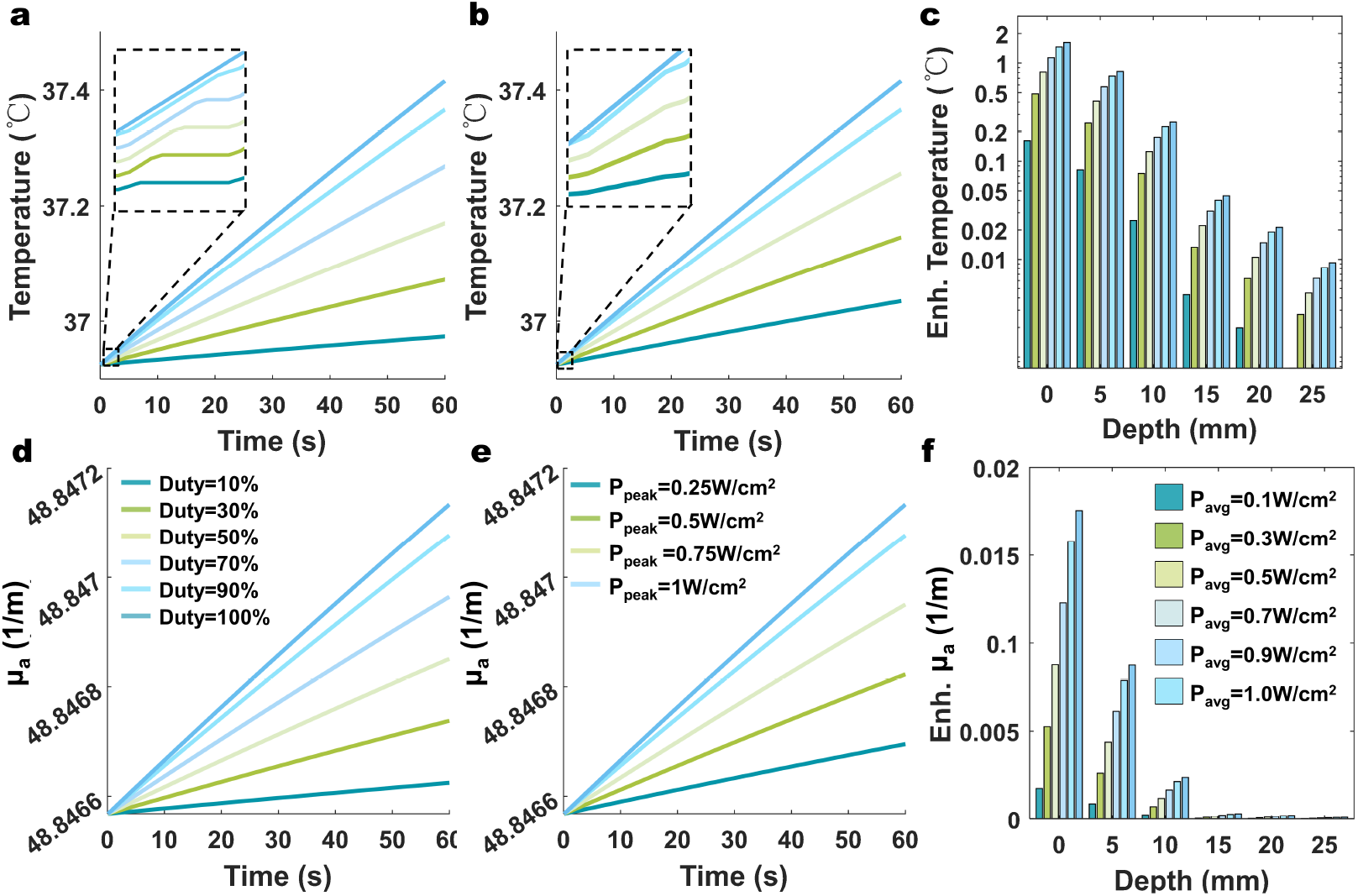
The thermal effects of tPBM. (a, d) Temporal evolution of temperature and absorption coefficient (*μ*_*a*_) across six duty cycles. (b, e) Temporal evolution of temperature and absorption coefficient under four pulsed peak power densities (P_peak_). (c, f) Depth profiles of the enhanced temperature and absorption coefficient under 5 average power densities (P_avg_) at t = 60 s. In the y-axis labels, Enh. denotes enhanced.

Fig. 4 further profiles the temperature-mediated regulation of light migration from perspectives of parameter sensitivity and spatiotemporal evolution in the realistic head model. Temporally, the photon flux exhibits a continuous reduction from approximately 410.465 to 410.365 W/m^2^ over the one-minute stimulation (Fig. 4a). A detailed view reveals the transient dynamic across the off-time gap: the photon flux exhibits no signs of recovery, and the onset flux of pulse N+1 remains marginally lower than the final flux of pulse N (Fig. 4a). Spatially, the depth-dependent decay of fluence (Fig. 4b) shows the photon absorption is largely confined to the superficial 0-10 mm. Specifically, following a 1-min stimulation, intracranial fluence (15 mm) drops to ∼0.1 J/cm^2^ at 1 W/cm^2^ (vs. ∼0.2 J/cm^2^ at the scalp) and to an upper quartile of ∼0.03 J/cm^2^ at 0.25 W/cm^2^. Furthermore, the fluence decays exponentially outward from the stimulation center (Fig. 4c), exhibiting a positively skewed distribution evidenced by the lowered medians (Fig. 4b). Parametrically, increasing the duty cycle from 10% to 100% reduces the flux by ∼0.1 W/m^2^ after a 1-min stimulation relative to the uncoupled model (Fig. 4a). Meanwhile, escalating the peak irradiance (P_peak_) from 0.25 to 1 W/cm^2^ boosts the upper quartile of intracranial fluence (15 mm) from 0.03 to 0.12 J/cm^2^. Concurrently, this higher optical intensity significantly widens the first-to-last pulse fluence gap from 0.002 to 0.186 J/m^2^ (Fig. 4c).

**Fig. 4.**
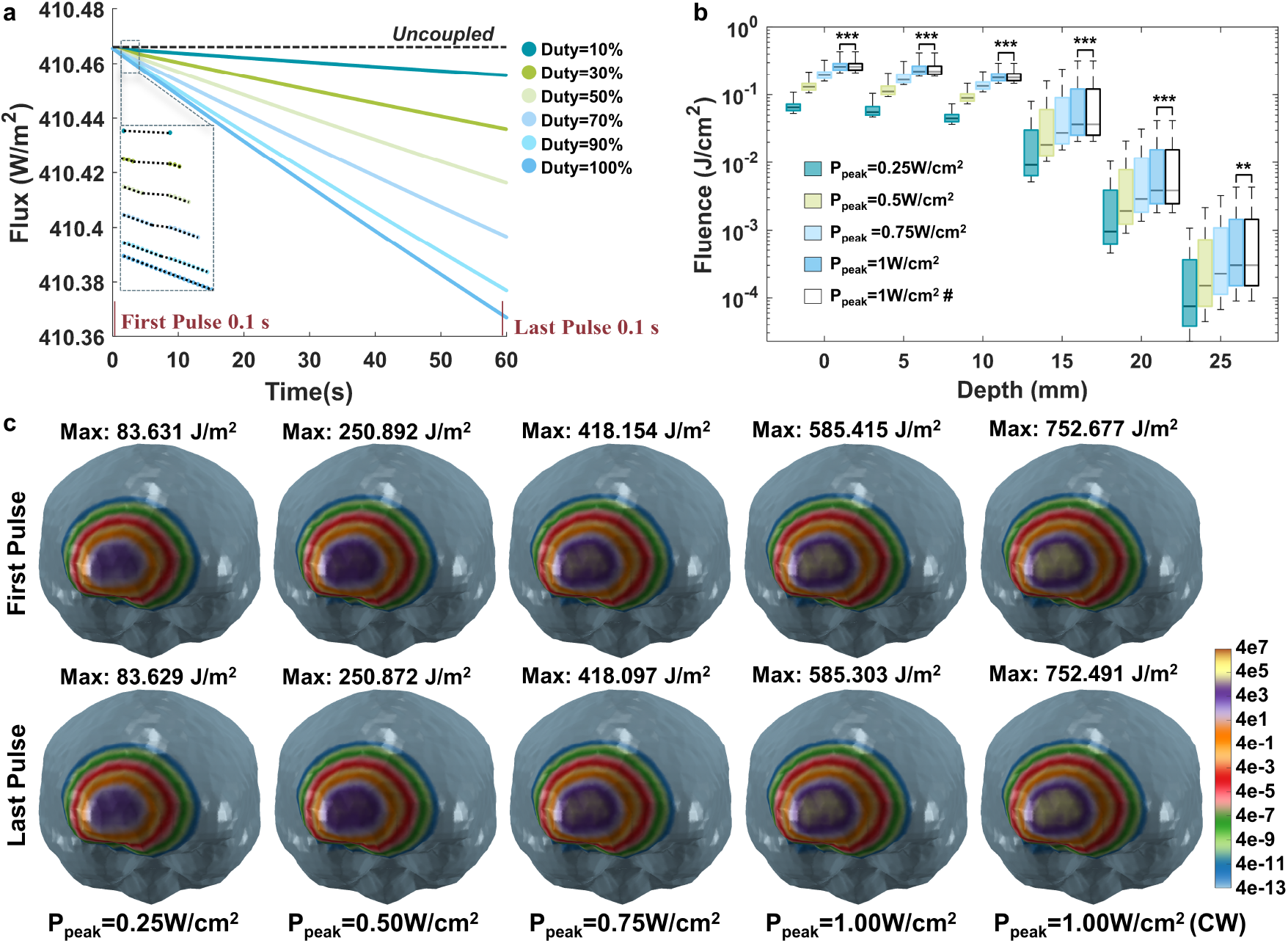
The thermally-modified light field. (a) Temporal evolution of photon flux across six duty cycles. (b) Depth profiles of 60-s cumulative photon fluence under four pulsed peak power densities (P_peak_). (c) Intracranial fluence distribution during the first versus last pulse cycles at four peak power densities (0.25-1 W/cm^2^, 90% duty cycle), compared with continuous-wave stimulation (1 W/cm^2^). Hash symbols (#) indicate results obtained by uncoupled model. Asterisks denote statistical significance based on two-tailed paired t-tests: * p < 0.05, ** p < 0.01, *** p < 0.001.

### 4.3. Comparison of Coupled and Uncoupled Model

To evaluate the necessity of bidirectional coupling, Fig. 5 compares the fluence and energy deposition predicted by the proposed coupled PTM against those from the conventional uncoupled model in the realistic head model. Overall, the coupled model consistently yields a lower fluence and a higher energy deposition (Fig. 5a, b, c). Temporally, the discrepancies in fluence and energy deposition between the two models widen exponentially, reaching 3 × 10^−4^ J/cm^2^ and 1.5 × 10^−4^ J/cm^3^ after one-minute stimulation (Fig. 5a). Spatially, the absolute differences in fluence and energy deposition decay exponentially with distance, falling below 2 × 10^−3^ J/cm^2^ and 1 J/cm^3^ beyond 10 mm, while the relative difference (the maximum percentage difference) in fluence increases slightly from the scalp (∼0.024%) to a depth of 15–20 mm (∼0.028%). Furthermore, the positively skewed distribution in deep-area boxplots and the spatial map (Fig. 5d) jointly confirm that significant discrepancies are highly localized at the irradiation site. Parametrically, higher duty cycles and greater P_peak_ amplify the deviations in both fluence (up to 3 J/m^2^) and maximal intracranial energy deposition (up to 3.217 J/m^3^) due to enhanced cumulative photothermal conversion. Similarly, increasing the optical intensity (P_peak_: 0.25 to 1 W/cm^2^, P_avg_: 0.1 to 1 W/cm^2^) elevates the absolute difference from 10^- 4^ to 10^-3^ J/cm^2^ and the relative difference from 0.001% to 0.01%.

**Fig. 5.**
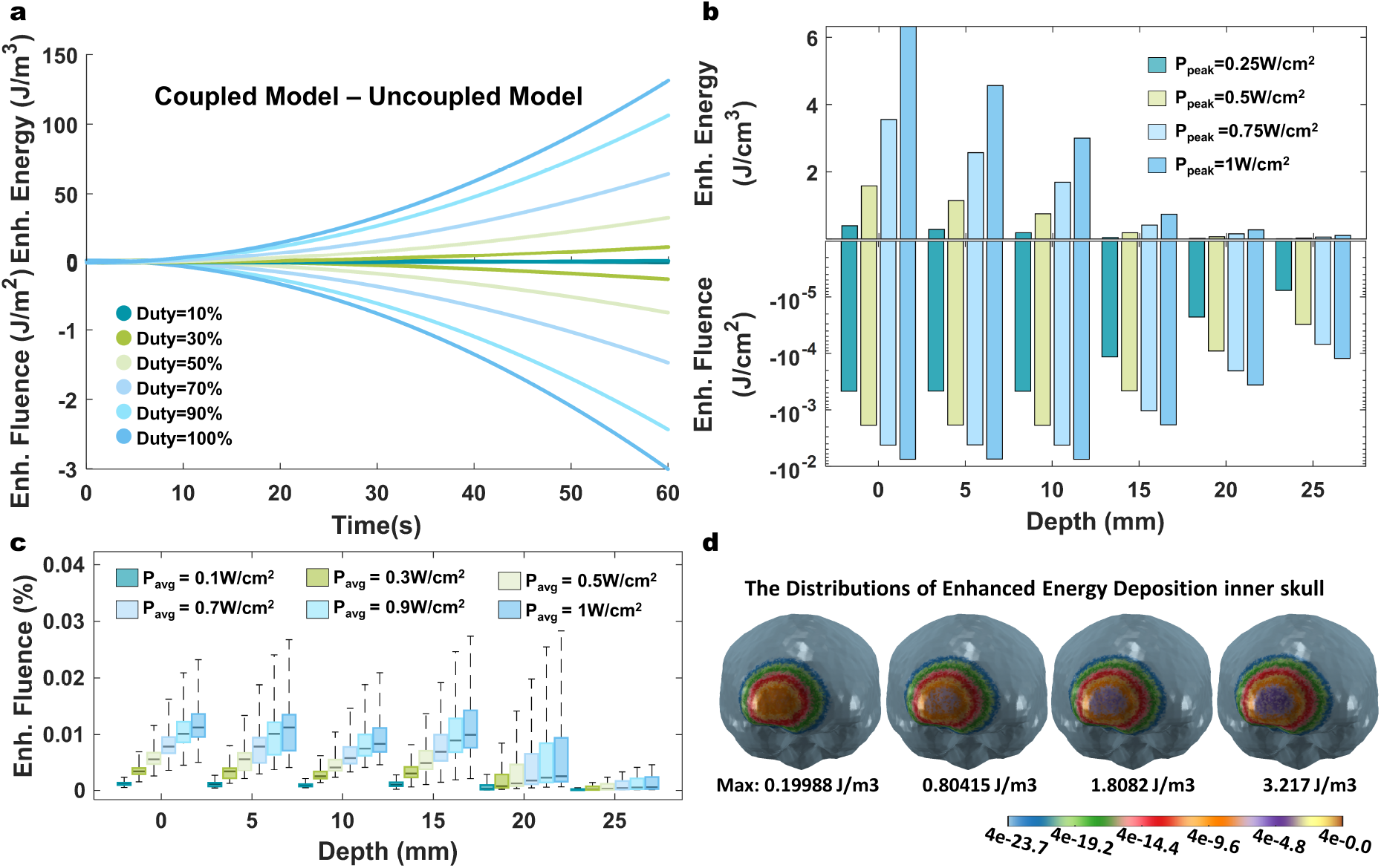
Comparison of the proposed photo-thermal bidirectional coupling model with the conventional uncoupled model. (a) Temporal evolution of energy deposition (top) and photon flux (bottom) across six duty cycles. (b) Depth profiles of 60-s enhanced energy deposition (top) and fluence (bottom) under four pulsed peak power densities (P_peak_). (c) Depth profiles of 60-s enhanced photon fluence under five average power densities (P_avg_). (d) Intracranial distributions of energy deposition under four P_peak_. In the y-axis labels, Enh. denotes enhanced.

### 4.4. Comparison of CW and PW

Fig. 6 compares the field distributions between CW and PW stimulation modes in the realistic head model. Capturing the transient spatiotemporal disparities between CW and PW modes underscores a unique advantage of the proposed PTM, as conventional static models inherently fail to resolve such dynamic photothermal variations. Macroscopically, the distribution differences (RDMs) of CW and PW for temperature, absorption coefficient, flux, and energy deposition across three tissues remain below 5 × 10^−7^ (Fig. 6a), indicating highly comparable overall field distributions. Notably, the maximal RDMs are strictly confined to the scalp, indicating that any discernible variations are predominantly superficial. Despite this overarching parity, a critical distinction emerges in localized penetration depth. Specifically, the PW mode achieves an enhanced penetration depth of approximately 1.43 mm over the CW mode, evaluated at an average power density of 0.5 W/cm^2^ and a flux threshold of 1 W/m^2^ (Fig. 6b). Furthermore, under identical energy density conditions, the PW mode exhibits a globally higher photon flux (fluence rate) across the spatial profile (Fig. 6c).

**Fig. 6.**
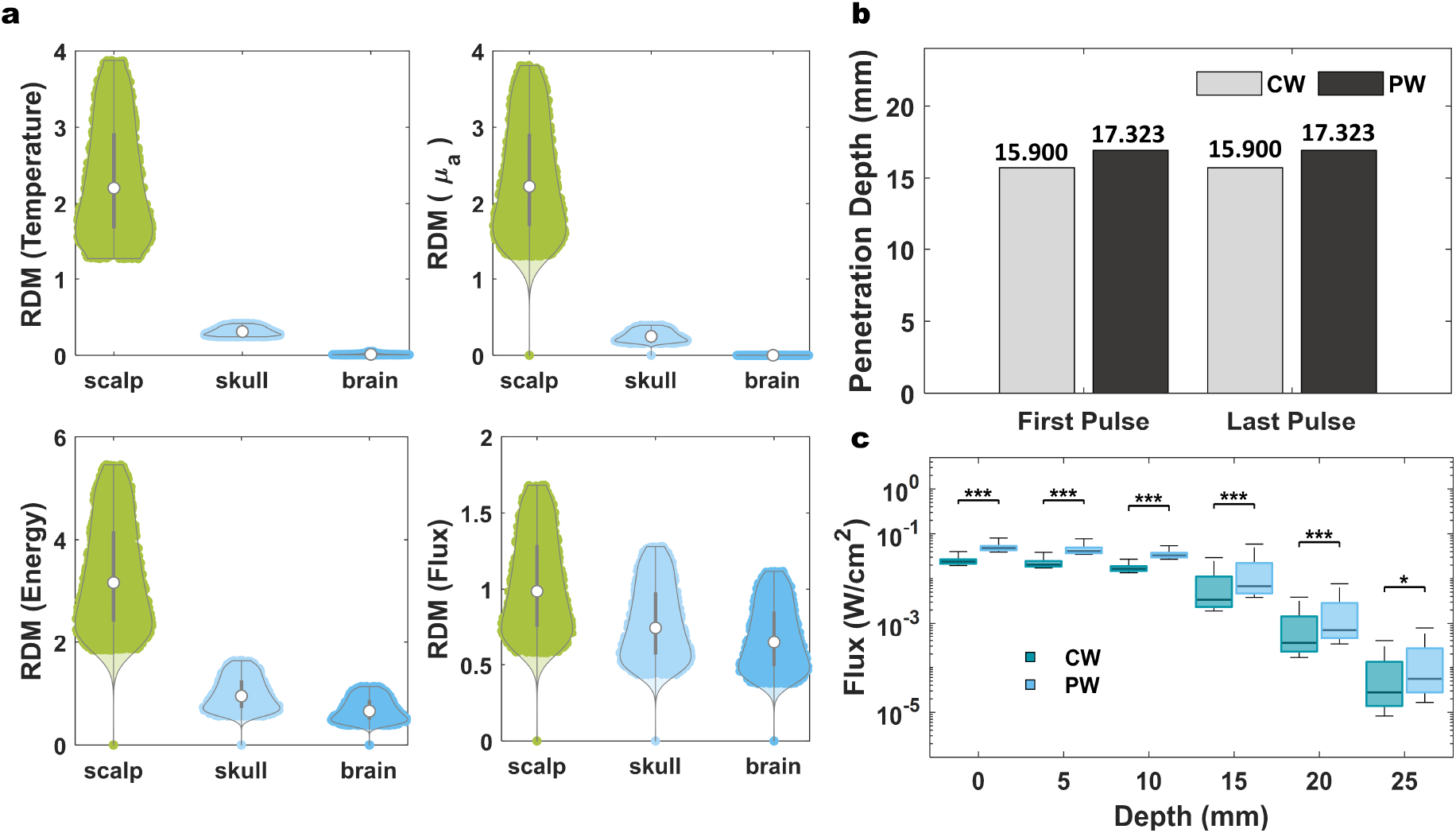
Comparison of continuous-wave (CW) and pulse-wave (PW) stimulations. (a) Relative difference measure (RDM, defined as the normalized distribution difference between CW and PW) for temperature, absorption coefficient (*μ*_*a*_), photon flux, and energy deposition at an average power density of 0.5 W/cm^2^ over 600 seconds. (b) Penetration depth for CW and PW, evaluated at the time points corresponding to the first and last pulses of the PW stimulation (pulse duration: 0.1 s per pulse; see Fig. 4a for the definition of the first and last pulses). (c) Depth profiles of photon flux for CW and PW at an average power density of 0.5 W/cm^2^ and a stimulation time of 599.95 s.

## 5. Discussions

The development of high-fidelity numerical models serves as the cornerstone for precise tPBM dose delivery. While current research predominantly attributes the primary effects of tPBM to photochemical interactions mediated by CCO absorption, the incident optical energy inevitably induces localized tissue heating [54, 55, 56, 57]. This thermal accumulation dynamically modulates tissue optical properties, thereby reshaping the optical field. To address this, this paper proposes a newly developed PTM that accounts for thermo-optical feedback by coupling the PBE with the RAM. The following sections evaluate the computational fidelity of this framework, analyze the spatial dosimetric trade-offs revealed by bidirectional coupling, and discuss its practical implications for parameter-dependent applicability and temporal modality optimization.

### 5.1. Validation and Spatiotemporal Dynamics of Bidirectional Coupling

To establish the computational reliability of the PTM, we benchmarked the coupled solver against an analytical solution [53]. The comparable dynamic ranges observed in the benchmark analysis confirm the numerical stability of the finite element formulation, ensuring the physical plausibility of the thermal bounds. Furthermore, the exponential decrease in RDMs is highly consistent with the intrinsic error-diffusion characteristics of parabolic partial differential equations governing thermal transport. Quantitatively, the persistent error reduction (2.35%–8.62%) and better boundary alignment in the coupled model validate its enhanced predictive accuracy. These improvements suggest that the bidirectional coupling mechanism actively suppresses boundary-induced computational errors, thereby providing a higher-fidelity replication of the exact analytical temperature field.

Simulations in the realistic head model reveal that tPBM dosing is an inherently non-static process [57]. This transient behavior is mediated by the RAM, which links localized tissue heating with a resultant rise in the absorption coefficient (*μ*_*a*_) [28, 29, 30]. Specifically, the strict spatiotemporal correspondence observed between thermal accumulation and the synchronous, quantitative escalation of *μ*_*a*_ effectively validates the bidirectional coupling mechanism established in the PTM. Fundamentally, this RAM-mediated thermal feedback induces a dynamic optical shading effect: the temperature-dependent rise in *μ*_*a*_ accelerates energy deposition while concurrently driving a continuous, nonlinear attenuation of photon fluence. By explicitly resolving this time-varying coupling, the PTM captures the critical temporal dynamics and spatial variation that stationary models inherently overlook. This nonlinear dynamic shading becomes particularly evident during pulsed stimulation. Rather than returning to baseline during the off-time phase, tissue temperature exhibits a transient overshoot followed by a plateau, indicating that the cessation of optical excitation does not correspond to an instantaneous thermal recovery [29, 58, 59]. Thus, subsequent pulses propagate through tissue with thermally preconditioned optical properties, leading to an onset fluence rate marginally lower than the terminal flux of the preceding pulse. Over a 60-s stimulation window, this cumulative thermal lag expands the fluence discrepancy between the initial and final pulses from 0.002 to 0.186 J/m^2^, as the elevated temperature locks *μ*_*a*_ at a higher state, exerting a persistent dampening effect on subsequent pulses.

Mechanistically, this non-monotonic off-time behavior is likely driven by delayed spatial heat redistribution, whereby residual heat accumulated in superficial high-absorption layers continues to diffuse toward deeper tissue after illumination ceases [23, 60, 61]. The subsequent plateau suggests incomplete thermal relaxation under the current pulse settings. Specifically, the 0.09-s off-time is insufficient to restore tissue temperature to the baseline, which serves as a plausible physical explanation for the limited discrepancy observed between PW and CW in later penetration simulations. It is worth noting that although significant thermal lag prevents full thermal recovery under current parameters, extending the off-time (e.g., via a lower stimulation frequency and duty cycle) may facilitate more thorough thermal relaxation. In essence, light propagation during tPBM emerges as a dynamically self-regulating process, where accumulated heat acts as a continuous negative feedback loop that attenuates the internal optical field over time.

### 5.2. Impact of Dynamic Optical Shifts on tPBM Dosimetric Evaluation

Comparative analysis between the PTM and conventional static (uncoupled) models reveals systematic dosimetric discrepancies. Over a 60-s stimulation, the static model yields a consistently higher photon fluence (by up to 3 × 10^−4^ J/cm^2^) but a lower energy deposition (by up to 1.5 × 10^−4^ J/cm^3^) relative to the dynamical PTM, a divergence that increases nonlinearly over time. Driven by the omission of thermal attenuation and the temperature-dependent escalation of *μ*_*a*_, this directional gap exposes a fundamental dual bias in static simulations: overestimating photon penetration while underestimating superficial energy confinement, aligning with prior reports [22]. This spatial dosimetric bias directly affects the calculation of required exposure durations, particularly when targeting specific energy thresholds (e.g., 0.9 to 15 J/cm^2^ for CCO activation [62]). To investigate whether such initial instantaneous deviation aggregate into therapeutically substantial discrepancies, we performed a longitudinal projection of these nonlinear dynamics. Under a 0.25 W/cm^2^ irradiance, our 60-s model yielded an intracranial fluence of ∼0.03 J/cm^2^ alongside an initial attenuation rate of 0.028%/min. Employing a linear extrapolation of this rate as a conservative estimate, we project that reaching the 0.9 J/cm^2^ lower threshold requires ∼31 min (cumulating ∼0.93 J/cm^2^), demanding ∼1 extra minute compared to static models; targeting the 15 J/cm^2^ upper threshold extends this gap significantly, requiring ∼40 additional minutes.

In summary, these results indicate that the effects of thermal feedback on tPBM simulation are both duration- and dosage-dependent. Under short-duration and low cumulative dose paradigms, its impact may remain relatively limited; however, under prolonged stimulation or high cumulative dose conditions, cumulative deviations introduced by dynamic optical-thermal coupling may become increasingly pronounced. Importantly, these deviations arise from the incomplete physical-mathematical representation of photo-thermal coupling in conventional formulations [48, 57, 63], underscoring their critical inadequacy for precise tPBM dosimetric evaluation, and therefore cannot be resolved solely through numerical refinement or algorithmic improvements.

### 5.3. Parameter Sensitivity and Model Applicability

This section further investigates parameter sensitivity and its implications for model applicability. For low-energy dosimetric planning involving short-duration exposures and low irradiation intensity, static uncoupled models offer a computationally efficient and sufficiently accurate approximation, circumventing the burden of dynamic field resolution. However, dosimetric discrepancies amplify substantially with prolonged optical energy input, such as extended on-times (higher duty cycles) and increased optical doses (higher peak power densities). Mechanistically, elevating the duty cycle amplifies energy input and diminishes the temporal window for thermal relaxation, while escalating the peak density magnifies instantaneous thermal transients, hindering baseline thermal recovery. Both parameters consequently drive higher local tissue temperatures [24], which exacerbate the temperature-dependent attenuation of the photon fluence, thereby underscoring the critical importance of the proposed PTM and its profound thermal modulation effect. Our simulations reveal that this effect produces a nonlinear increase in the discrepancy between coupled and uncoupled predictions, highlighting the necessity of the proposed PTM. This suggests that, while our earlier conservative linear projection provides a lower−bound baseline for long−term error accumulation, the true error is likely larger, and capturing the accelerated energy dynamics of tPBM across different intensities indeed necessitates a nonlinear coupled model. Additionally, spatial analyses demonstrate that the most pronounced deviations co-localize with the focal target region; thus, relying on spatially averaged error metrics may underestimate actual therapeutic discrepancies. Consequently, while static models remain viable for low-intensity paradigms (e.g., P_peak_ ⩽ 1 W/cm^2^ for ⩽ 1 min) with relaxed precision constraints, bidirectional coupling becomes indispensable as energy delivery and dosimetric strictness escalate (e.g., 15 J/cm^2^ CCO activation). Ultimately, these findings demonstrate that tPBM induces a dose-dependent, time-accumulative, and superficially concentrated thermal effect that continuously reshapes the local optical absorption profile.

### 5.4. Modality Selection: A Field-Level Perspective on PW and CW

A primary challenge in selecting optimal stimulation modes for tPBM is elucidating the divergent biophysical mechanisms underlying PW and CW irradiation. Although empirical differences are documented, direct physical explanations remain scarce, often relying on indirect hypotheses such as neural entrainment [64, 65]. This mechanistic gap persists because current uncoupled models lack dynamic spatiotemporal resolution. When comparing CW and PW modalities under identical average energy input, our simulations reveal highly comparable macroscopic field distributions, likely attributable to the short thermal relaxation time. Despite this bulk similarity, PW irradiation achieves a localized advantage, extending penetration depth by approximately 1.43 mm directly beneath the irradiation center. Optically, this deeper penetration is a direct consequence of the higher peak irradiance required by PW protocols to maintain equivalent average power. This higher peak irradiance also corresponds to a globally elevated photon flux under identical energy density (Fig. 6). While conventional tPBM literature predominantly posits that neuromodulatory effects are strictly energy-dependent, emerging evidence suggests irradiance acts as a binary gating switch for modality-specific outcomes, where stronger peak irradiance (i.e., higher photon flux) in PW protocols may trigger distinct physiological responses [66]. Overall, the proposed coupled framework could offer a complementary, physical-field perspective for optimizing temporal patterns in neuromodulation.

### 5.5. Future Direction

Although the PTM establishes a dedicated framework for bidirectional photothermal dosimetry, three primary limitations warrant acknowledgment. First, the current simulations utilize a simplified three-layer head model. Future studies could enhance anatomical fidelity by transitioning to a more complex six-layer vascularized model. Second, resolving these coupled multi-physics fields imposes a severe computational bottleneck. To ensure stability, the current solver demands a rigid 0.01-second timestep, making simulation of longer durations (>10 min) computationally prohibitive, with an estimated runtime of 100 days on standard hardware. Future studies could employ adaptive time-stepping algorithms and GPU acceleration to make long-duration tPBM simulations practically feasible. Third, the current validation is limited to analytical benchmarks, and experimental validation of the coupled framework is still lacking. Future studies should validate model predictions using tissue-mimicking phantoms, ex vivo preparations, or in vivo animal models, followed by clinical studies to assess the practical utility of the proposed framework for tPBM protocol optimization.

## 6. Conclusion

This study develops a novel PTM to simulate the photo-thermal spatiotemporal dynamics of tPBM. The model addresses the fundamental limitations of current uncoupled approaches by bidirectionally coupling the light and heat fields through the integration of the PBE and the RAM. By ensuring strict energy conservation, the PTM yields physically rigorous predictions of attenuated photon fluence and concurrently elevated localized energy deposition, thereby correcting the systematic overestimation of fluence and underestimation of tissue temperature inherent in uncoupled models. Analytic validation confirms the accuracy of the solution for the coupled equations. Comparative simulations demonstrate that the predictive deviation between the PTM and uncoupled models amplifies nonlinearly with increasing exposure duration and intensity, underscoring the necessity of coupled modeling for high-dose or prolonged protocols. Furthermore, a key finding enabled by this framework is the identification of a distinct physical-field disparity between the PW and CW irradiation; specifically, PW achieves a 1.43 mm greater penetration depth under equivalent average power. Ultimately, the PTM provides an integrative framework that unifies the optical and thermal physics of tPBM, establishing a robust foundation for precise dosimetric evaluation and serving as a vital tool for future mechanistic and clinical investigations.

## 7. CRediT authorship contribution statement

**Ting Zhang:** Writing – review & editing, Writing – original draft, Conceptualization, Methodology design, Formal Analysis, Data Curation. **Yutao Chen:** Formal Analysis. **Xin Zeng:** Data Curation. **Ge Zhang:** Writing – review & editing. **Feiyan Wang:** Writing – review & editing. **Daqing Guo:** Writing – review & editing, Conceptualization, Supervision, Funding acquisition. **Dezhong Yao:** Writing – review & editing, Conceptualization, Supervision, Funding acquisition.

## 9. Acknowledgements

We sincerely thank Dr. Hakan Erkol (Boğaziçi University) for his generous guidance on analytical modeling, and Dr. Xiaoping Li (University of Electronic Science and Technology of China) for his valuable assistance with finite element numerical methods. Their insightful suggestions greatly improved this work.

This work was supported in part by the Brain Science and Brain-like Intelligence Technology-National Science and Technology Major Project under grant 2022ZD0208500, in part by the Lingang Laboratory under grant LGL-1987, in part by the National Key Research and Development Program of China under grant 2023YFF1204200, in part by the Sichuan Science and Technology Program under grant 2024NSFJQ0004, grant DQ202410 and grant 2024NSFTD0032.

## Supporting Information Text

Detailed finite element derivation process of the light diffusion and Pennes bioheat equations is provided here.

## S1. Finite Element Analysis for Light Diffusion Equation

This section including three parts: weakening, discretization, and solving. The diffusion approximation of PBM [1, 2] is

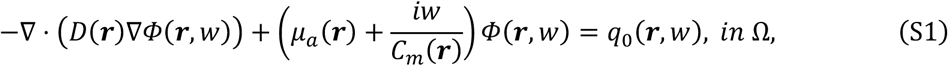

where Ω is the solution domain of the target light field, i.e., biological tissue; *Φ*(***r***, *w*) is the photon flux at position ***r***, and modulation frequency *w*, with unit 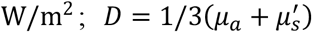 is the diffusion coefficient; *μ*_*a*_ and 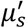 are the absorption and reduced scattering coefficients, respectively; *C*_*m*_(***r***) is the light speed in the biological tissue at position ***r***; *q*_0_(***r***, *w*) is the isolated source at position ***r***, and modulation frequency *w*.

The corresponding air boundary is depicted by Robin boundary Condition.

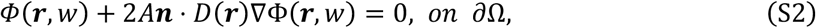

where ∂Ω is the external surface of Ω and ***r*** is a point on ∂Ω; ***n*** is the outward-pointing normal, *A* depends on the refractive index (RI) mismatch between scalp and air.

### S1.1 Weakening

A weak formulation of equation (S1) can be obtained using integration by parts after multiplied by *φ* ∈ ℍ^1^(Ω)

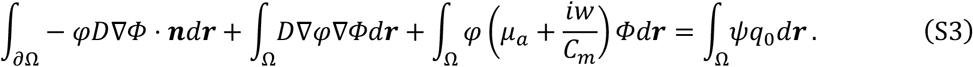

Applying the boundary condition (S2) to formula (S3), we get the weak formulation of light diffusion equation (LDE)

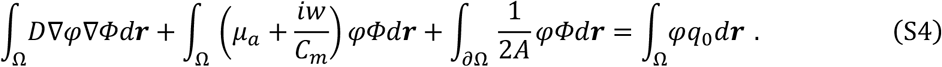

### S1.2 Discretization

The discretization scheme depends on the mesh types adopted for representing the head domain, with different types corresponding to distinct shape functions. For finer characterization of complex geometrical boundaries, a tetrahedral mesh is employed for discretization. Given a mesh parameter *h*, let Ω_*h*_ represent the discrete domain and *N* the number of mesh nodes. The approximate photon flux Φ_*h*_ ∈ Ω_*h*_ admits the representation.

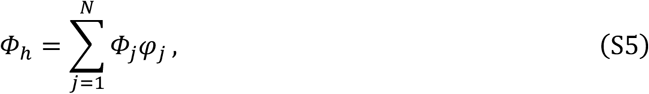

where *Φ*_*j*_ is the photon flux of the mesh node *j*; *ψ*_*j*_ is the linear shape function in three-dimensional space. Then, apply the weak formulation (S4) to the discrete domain Ω_*h*_, we get

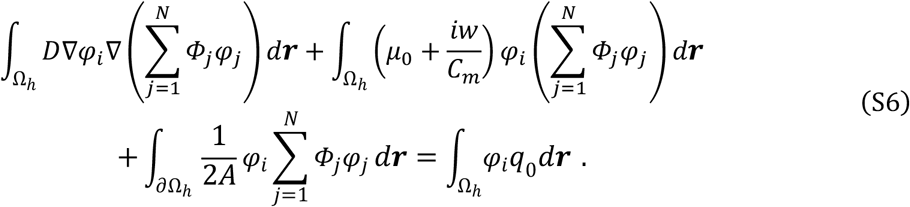

Interchanging the order of the integral and summation yields the finite element formulation of the LDE.

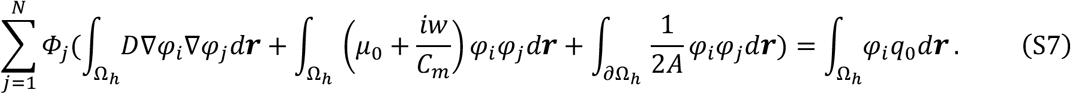

### S1.3 Solving

Rearranging the integral equation above leads to the following system of linear equations.

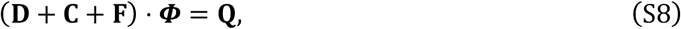

where **×, C, F** ∈ ℝ^*N*×*N*^ are the terms related to the geometry and the physical properties of tissues; **Q** ∈ ℝ^*N*×1^ is a vector related to light sources; ***Φ*** = (*Φ*_1_, *Φ*_2_, …, *Φ*_*N*_) ∈ ℝ^*N*×1^ is the numerical solution of this system provides the desired approximation. A complete derivation is available in the Supplementary Material. **×, C, F** and **q**_0_ are given by:

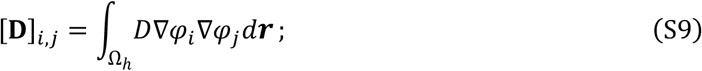

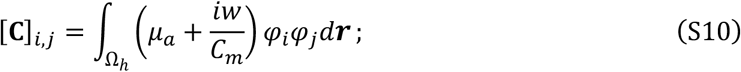

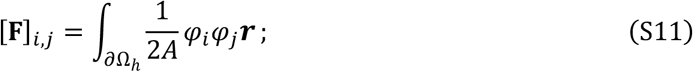

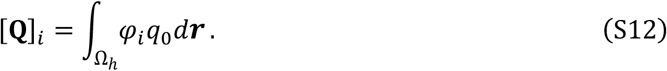

The photon flux **Φ** can be obtained by solving the aforementioned linear system.

## S2. Finite Element Analysis for Bioheat Equation

This section’s derivation proceeds analogously to Section I, with the only distinctions occurring in the details stemming from the specific form of the governing equation. The bioheat transfer [3] is modeled by the following Pennes Bioheat Equation (PBE):

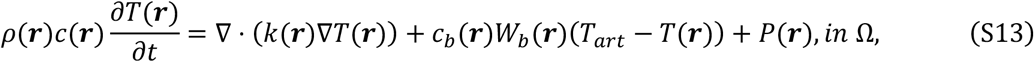

where Ω is the solution domain of the target thermal field, i.e., head tissue; *T*(***r***) is the temperature of biological tissue; *ρ*(***r***), *c*(***r***) and *k*(***r***) represent the tissue density, specific heat, and thermal conductivity respectively; *T*_*art*_ signifies constant arterial temperature (set at 37 °C in this paper); *w*_*b*_(***r***) and *c*_*b*_(***r***) are the perfusion rate and specific heat of blood respectively; *c*_*b*_(***r***)*W*_*b*_(***r***)(*T*_*art*_ − *T*(***r***)) represents the thermal obtained from the blood perfusion; *P*(***r***) describes the energy received from external sources, unit: W/m^3^.

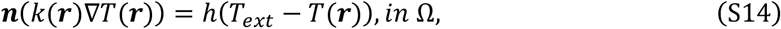

where ∂Ω is the external surface of Ω and ***r*** is a point on ∂Ω; ***n*** is the outward-pointing normal; *h* is the heat transfer coefficient, assigned a value of 6.2 W ⋅ *m*^−2^ ⋅ K^−1^ [4], and *T*_*ext*_ is the ambient temperature, set to 25 °C.

### S2.1 Weakening

A weak formulation of formula (S13) can be obtained using integration by parts after multiplied by *φ* ∈ ℍ^1^(Ω)

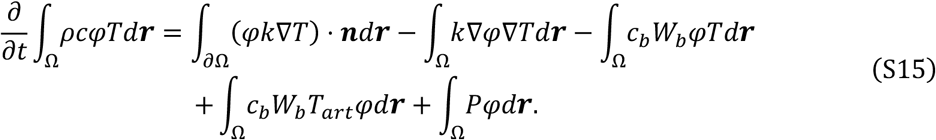

Applying the boundary condition (S14) to (S15), we get the final weak formulation of PBE

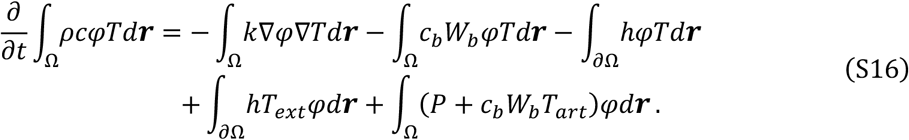

### S2.2 Discretization

For the defined discrete domain Ω_*h*_ (a tetrahedral mesh with parameter *h*), the temperature approximation *T*_*h*_ ∈ Ω_*h*_ is expressed as

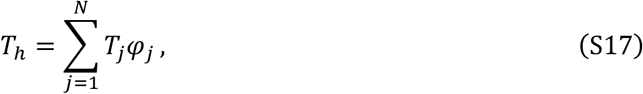

where *φ*_*j*_ is the temperature of the mesh node *j*; *ψ*_*j*_ is the linear shape function in three-dimensional space.

Then, applying the weak formulation (S16) to the discrete domain Ω_*h*_, we get

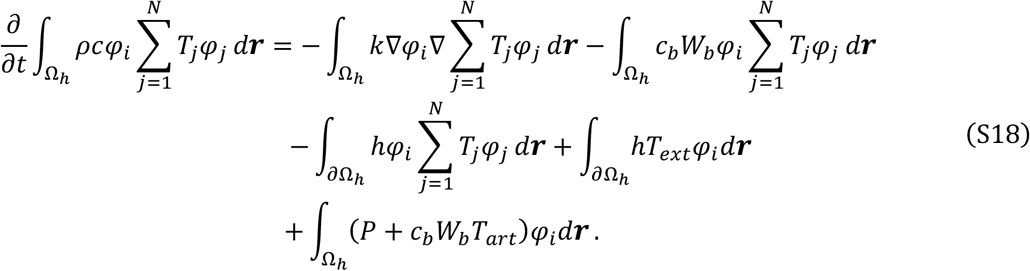

By interchanging the integral and summation operators, we arrive at the finite element formulation of PBE.

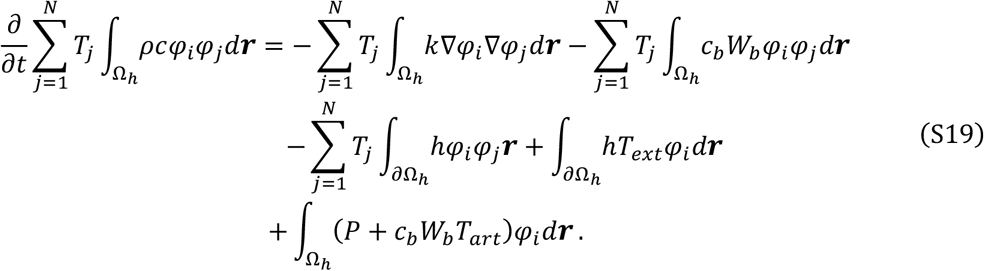

### S2.3 Solving

By organizing the above integral equation, we obtain the following linear system of equations.

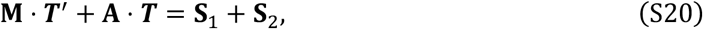

where **M, A** ∈ ℝ^*N*×*N*^ denote the mass and stiffness matrices; the vectors **S**_1_, **S**_2_ ∈ ℝ^*N*×1^ represent source terms, with **S**_1_ incorporating exchange with blood and the environment, and **S**_2_ due to the light absorption; and ***T*** = (*T*_1_, *T*_2_, …, *T*_*N*_) ∈ ℝ^*N*×*N*^ is the discrete solution vector for temperature. The complete derivation is available in the Supplementary Material. The explicit expression for **M, A, S**_**1**_, **S**_**2**_ are defined as follows:

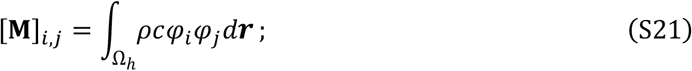

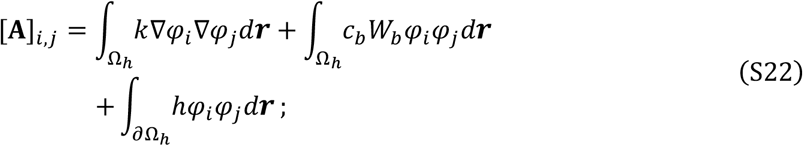

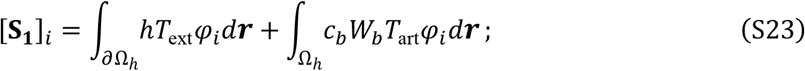

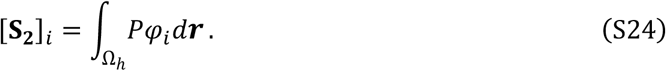

By solving the above linear system, we obtain the spatial distribution of temperature ***T***.

## S3. Analytical Expressions for Photon Density and Temperature Rise

The analytical solution used for numerical validation in the main text (Section 4.1) is taken from Erkol et al. (2020) [5]. This solution describes the steady-state photon density and the subsequent time-dependent temperature rise induced by continuous-wave laser heating in a two-layer cylindrical turbid medium, under the diffusion approximation to the radiative transfer equation and the Pennes bioheat equation.

### S3.1. Photon Flux

The photon density Φ(*r*, ϕ) [W/mm^2^] in the two-layer cylindrical medium (inner radius *R*_in_, outer radius *R*) is given by:

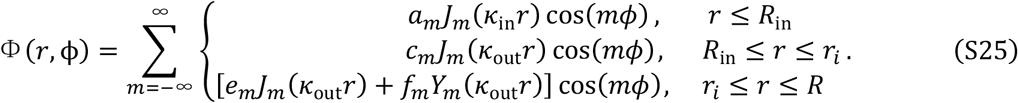

Here, *J*_*m*_ and *Y*_*m*_ are the Bessel functions of the first and second kinds, respectively; 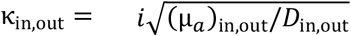 is the absorption coefficient; 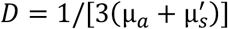 is the diffusion coefficient, with 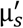 being the reduced scattering coefficient; *r* is the radial coordinate; ϕ is the angular coordinate; cos(*m*ϕ) is the angular eigenfunction arising from the separation of variables in polar coordinates, with the cosine term retained due to the symmetry about the *x*-axis; 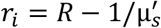 is the position of the isotropic point source; and *m* is the azimuthal mode number. In this work, the infinite series over $m$ is truncated at |*m*| = 30.

The coefficients *a*_*m*_, *c*_*m*_, *e*_*m*_, and *f*_*m*_ are determined by the interface continuity conditions at *r* = *R*_*i*n_, the source discontinuity at *r* = *r*_*i*_, and the Robin boundary condition at *r* = *R*. Their explicit expressions are provided in Eqs. (19) - (21) of Erkol et al. (2020) [5].

### S3.2. Temperature

The time-dependent temperature rise Δ*T*(*r*, ϕ, *t*) = *T*(*r*, ϕ, *t*) − *T*_s_ [K], where *T*_s_ is the ambient temperature, is given by:

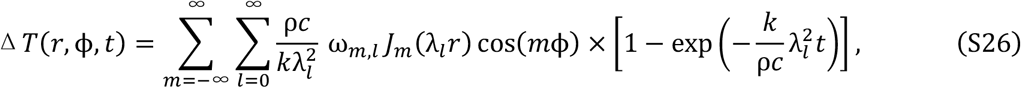

where ρ is the medium density [g/mm^3^], *c* is the specific heat [J/(g·K)], *k* is the thermal conductivity [W/(mm·K)], and λ_*l*_ are the eigenvalues determined by the convective boundary condition at *r* = *R*:

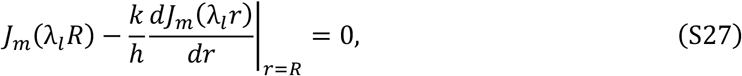

with *h* being the heat transfer coefficient [W/(mm^2^·K)]; *l* is the radial mode number. The coefficients ω_*m*,*l*_ are obtained by projecting the heat source term μ_*a*_Φ(*r*, ϕ)/(ρ*c*) onto the eigenfunctions J_*m*_(λ_l_ r) cos(*m*ϕ); their detailed expressions are given in Eqs. (36) - (41) of Erkol et al. (2020) [5].

### S3.3. Parameters Used in This Work

The specific geometrical and optical parameters adopted for the numerical validation in this study are listed in Table S1.

**Table S1.** Geometrical, optical, and thermal parameters used for the numerical validation.

| Parameter | Symbol | Value |
| --- | --- | --- |
| Outer radius | $R$ | 25 mm |
| Inner (inclusion) radius | $R_{\text{in}}$ | 18 mm |
| Phantom height | $H$ | 1100 mm |
| Source position (radial) | $r_i$ | $R - 1/\mu'_s$ |
| Source angle | $\phi_i$ | 0° (along x-axis) |
| Source arc | --- | 60° |
| Background absorption coefficient. | $\mu_a^{\text{out}}$ | 0.028 mm<sup>-1</sup> |
| Inclusion absorption coefficient. | $\mu_a^{\text{in}}$ | 0.060 mm<sup>-1</sup> |
| Background reduced scattering coefficient. | $\mu'_s$ | 2.050 mm<sup>-1</sup> |
| Inclusion reduced scattering coefficient. | $\mu'_s$ | 2.872 mm<sup>-1</sup> |
| Thermal conductivity | $k$ | $3.42 \times 10^{-4}$ (W · K<sup>-1</sup>mm<sup>-1</sup>) |
| Density | $\rho$ | 0.0011 kg/mm<sup>3</sup> |
| Specific heat | $c$ | 3.662 (J · g<sup>-1</sup>K<sup>-1</sup>) |
| Heat transfer coefficient | $h$ | 6.2 × 10<sup>-3</sup> (W · K<sup>-1</sup>mm<sup>-1</sup>) |
| Ambient temperature | $T_s$ | 25 °C |

Note: For the detailed derivation process of the above analytical solutions (including the integral Green's function method and the separation of variables technique), readers are referred to the original publication: Erkol et al. (2020) [4].

